# Mechanisms of KCNQ1 Channel Dysfunction in Long QT Syndrome Involving Voltage Sensor Domain Mutations

**DOI:** 10.1101/231845

**Authors:** Hui Huang, Georg Kuenze, Jarrod A. Smith, Keenan C. Taylor, Amanda M. Duran, Arina Hadziselimovic, Jens Meiler, Carlos G. Vanoye, Alfred L. George, Charles R. Sanders

## Abstract

Loss-of-function (LOF) mutations in human *KCNQ1* are responsible for susceptibility to a life-threatening heart rhythm disorder, the congenital long-QT syndrome (LQTS). Hundreds of *KCNQ1* mutations have been identified, but the molecular mechanisms responsible for impaired function are poorly understood. Here, we investigated the impact of 51 KCNQ1 variants located within the voltage sensor domain (VSD), with an emphasis on elucidating effects on cell surface expression, protein folding and structure. For each variant, the efficiency of trafficking to the plasma membrane, the impact of proteasome inhibition, and protein stability were assayed. The results of these experiments, combined with channel functional data, provided the basis for classifying each mutation into one of 6 mechanistic categories. More than half of the KCNQ1 LOF mutations destabilize the structure of the VSD, resulting in mistrafficking and degradation by the proteasome, an observation that underscores the growing appreciation that mutation-induced destabilization of membrane proteins may be a common human disease mechanism. Finally, we observed that 5 of the folding-defective LQTS mutants are located in the VSD S0 helix, where they interact with a number of other LOF mutation sites in other segments of the VSD. These observations reveal a critical role for the S0 helix as a central scaffold to help organize and stabilize the KCNQ1 VSD and, most likely, the corresponding domain of many other ion channels.

**One Sentence Summary:** Long QT syndrome-associated mutations in KCNQ1 most often destabilize the protein, leading to mistrafficking and degradation.

## Introduction

The KCNQ1 (Kv7.1) voltage-gated potassium channel contributes to a variety of physiological processes, most notably when it complexes with the KCNE1 accessory protein to generate the slow delayed rectifier current (Iks) of the cardiac action potential (*1-6*). Heritable mutations in *KCNQ1* resulting in channel dysfunction lead to type 1 long QT syndrome (LQTS), a potentially life-threatening predisposition to cardiac arrhythmia (*6, 7*). However, many patients with LQTS are unaware of their condition.

More than 600 KCNQ1 mutations associated with LQTS have been identified and this number continues to grow (*8-10*). While progress in functional characterization of LQTS-associated KCNQ1 mutations has been made (*11-16*), the mechanistic basis of channel dysfunction for most mutations is not known. Moreover, expanding use of exome/genome sequencing contributes to a growing database of unclassified human KCNQ1 mutations. These “variants of unknown significance” (VUS) present a medical quandary in the context of genetic testing for LQTS and the medical decision of who to preemptively treat with invasive procedures such as implantation of a defibrillator (*17*). The dearth of direct experimental data to determine how mutations alter channel function has prompted bioinformatic and modeling efforts to predict pathogenicity (*16, 18-20*). However, results from these *in silico* approaches are not considered strong evidence by genetic testing standards established by the American College of Medical Genetics and Genomics (ACMG) (*21*). This is in contrast to *in vitro* experimental data documenting mutation-induced loss of function for a disease-linked protein, which the ACMG guidelines categorize as strong evidence for pathogenicity (*21*).

Here we assess medium throughput *experimental* studies of KCNQ1 variants as a route to assessing their functional and biochemical consequences and determining the mechanistic basis for pathogenicity. A set of 51 single site mutants affecting sites located in the KCNQ1 voltage sensor domain (VSD) were compared to the wild type (WT) channel. A cross-section of mutants was selected to represent three classes of variants: known LQTS mutations, documented human VUS, and mutations predicted to be neutral based on WT occurrence within non-human KCNQ1 orthologs. The results of this work implicate the underlying mechanisms for LQTS-associated loss-of-function (LOF) mutations and also support the notion that most of the examined VUS are deleterious. Moreover, analysis of mutations that impact amino acid sites either located in the VSD S0 segment or that directly interact with S0 provides evidence that this overlooked structural element is critical for channel stability and function.

## Results

### Structure mapping of 51 human KCNQ1 variants

The KCNQ1 mutations chosen for study in this study represent three groups: (i) 17 previously associated with LQTS, (ii) 21 human VUS for which there has been insufficient data for classification as benign or pathogenic, and (iii) 13 designed variants predicted to be benign. The predicted benign variants were based on swapping in an amino acid observed to be divergent from the WT sequence in a non-human ortholog of KCNQ1 (*22*). For example, site 104 in human KCNQ1 is a threonine but is serine at the corresponding site in chameleon and in finch KCNQ1. The T104S mutation is therefore predicted to exhibit no significant functional consequences. All mutations are in the VSD and include many known single residue LQTS mutations in the S0 and S1 segments (fig. S1A). Figures S1B and S1C show their locations in the recently-determined structure of the voltage sensor of *Xenopus laevis* KCNQ1 (*23*).While the present study was carried out using human KCNQ1, high homology indicates the *Xenopus* structure can be assumed to be an excellent model for human KCNQ1. As show in fig. S1B, most of the mutations examined in this work affect the cytosol-facing portion of the VSD, although mutations in other parts of the VSD are also included.

As described elsewhere (*22*) and summarized in table S1 each of the 51 mutants was co-expressed in CHO-K1 cells with WT human KCNE1 and functionally characterized using whole cell electrophysiology (EP) to assess the effect of each mutation on Iks amplitude, voltage-dependence of activation and deactivation kinetics. The EP data identified 32 KCNQ1 mutants that failed to yield at least 65% of peak WT K^+^ channel current and were therefore classified as LOF. The 65% value chosen for the cutoff is based on the approximate boundary between the range of currents observed for known LQTS mutants and benign mutants based on an extensive literature review (see also (*16*)). The underlying mechanisms for LOF in these 32 mutants were not established by the functional data alone, leading to the experiments of the present study.

### Mistrafficking is a common cause of mutation-induced channel LOF

We developed a method for quantitating KCNQ1 cell surface and total expression. Using a fully functional form of KCNQ1 in which a Myc epitope is inserted into the extracellular loop connecting S1 and S2 (*24*) we transiently expressed WT and variant channels in HEK293 cells followed by labeling the cell surface population of KCNQ1 with an anti-Myc antibody conjugated to phycoerythrin (PE). The cells were then permeabilized and the intracellular population of KCNQ1 was labeled with a second anti-Myc antibody, this time conjugated with Alexa Fluor-647. The cells were then subjected to flow cytometry to measure the single cell intensities of PE and Alexa Fluor-647 fluorescence, providing quantification of both total and surface channel expression levels (fig. S2). Fig. 1A shows the quantified cell surface levels for KCNQ1 mutants and reveals that many LQTS and VUS mutants exhibit much lower surface expression than WT. In considering these results, we acknowledge that HEK293 cells are an imperfect replacement for cardiomyocytes or CHO-K1 cells (optimal for functional studies of KCNQ1). However, the use of cardiomyocytes for high/medium throughput EP and trafficking experiments of KCNQ1 mutants co-transfected with KCNE1 is not currently feasible. Moreover, we found that HEK293 cells yielded higher signal-to-noise measurements in our trafficking assays than CHO-K1. Fortunately, as will be shown the use of HEK293 cells yielded results are that are strikingly consistent with what is expected based both on the previous EP results, disease classification, and predictions for this panel of 51 KCNQ1 variants.

**Figure 1.**
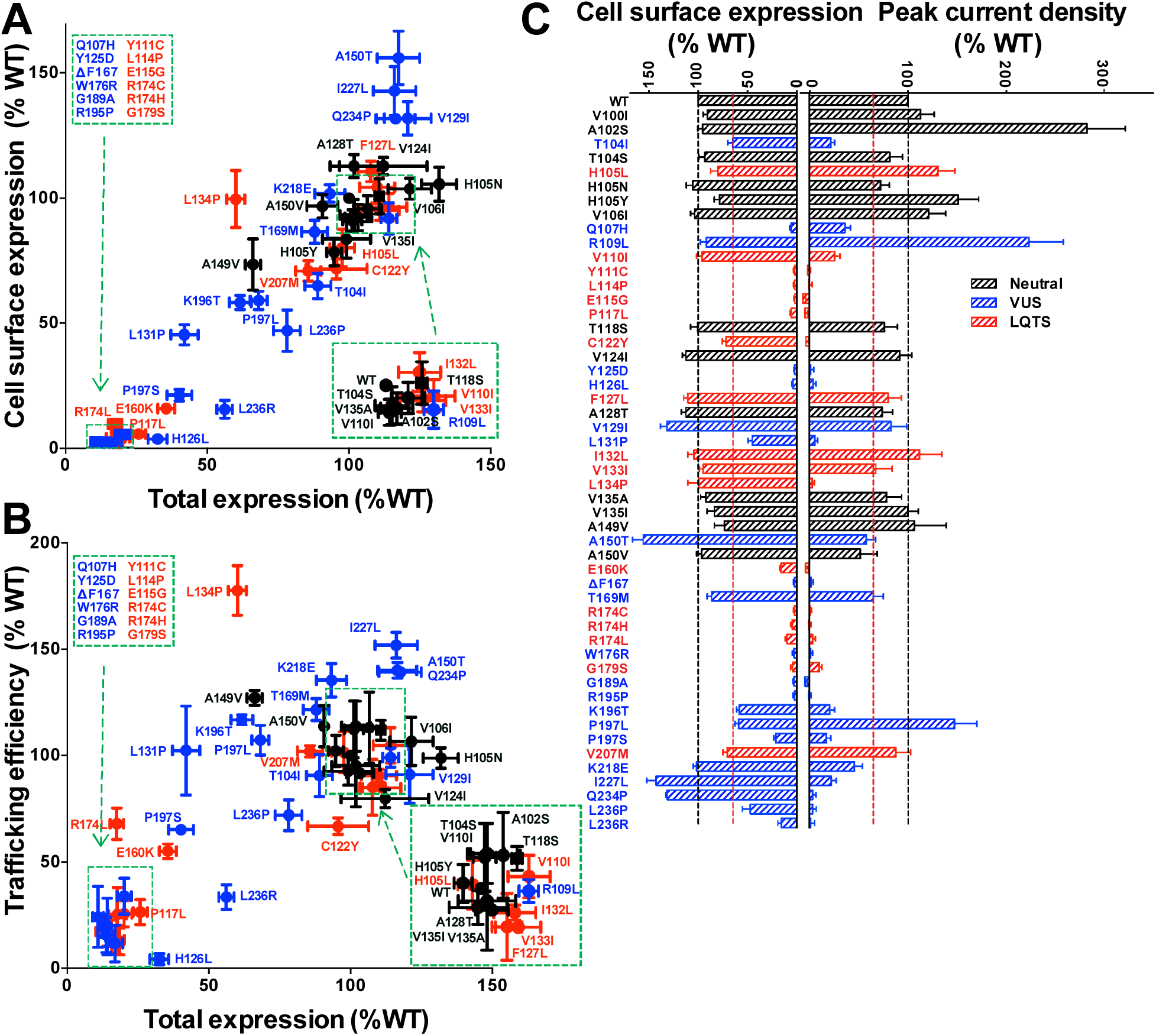
KCNQ1 expression levels and surface-trafficking efficiencies. Data are colored coded: LQTS mutant (red) VUS (blue) or predicted neutral polymorphism (black). For all three panels reported expression levels are relative to WT results and are expressed as mean ± S.E.M. based on least three independent experiments. (**A**) Total expression levels plotted as a scatter plot versus surface expression. The inset boxes illustrate which sets of mutants yield near-0 or WT-like (100%) surface expression levels. Data labeled with an asterisk indicate those for which the measured KCNQ1 surface protein level was statistically different from the WT level (P<0.05). (**B**) Surface trafficking efficiency versus total expression level, where efficiency is defined as: [(surface)_mutant_/(total)_mutant_]/[(surface)_WT_/(total)_WT_] X 100. (**C**) Cell surface expression levels as measured in this study versus K^+^ channel peak current density, as originally reported elsewhere(*22*). The vertical red lines indicate 65% of WT, the effective upper limit cut-off for LOF.

Fig. 1A also shows that there is a strong correlation between cell surface expression levels and total (surface + internal) expression. This correlation can be explained based on two factors. First, mutant KCNQ1 channels exhibit lower cell surface expression levels because of reduced total expression. Secondly, for many mutants exhibiting low total expression it was seen that surface expression is even lower than predicted based on the degree of reduction seen for total expression. A replot of the data is shown in Fig. 1B that illustrates the relative contributions of these two factors: this panel shows the surface trafficking *efficiency* versus total expression, where efficiency is defined as [cell surface]/[total] expression for the mutant relative to [cell surface]/[total] expression for WT (× 100). It is seen that 18 of the 23 mutants exhibiting ≤65% of total WT expression also exhibit cell surface trafficking *efficiencies* that are ≤65% that of WT. This indicates that not only do some mutants exhibit low total expression, but also that in such cases the modest population of channel that does express is surface trafficking-impaired. We note the gratifying result that all 13 predicted benign mutants exhibited both total and surface expression levels that are similar to WT.

Figure 1C and table S1 present peak channel current densities for each mutant (*22*) juxtaposed with the cell surface expression levels. This reveals that channel function and surface trafficking are often strongly coupled, indicating that reduced levels of KCNQ1 at the plasma membrane are the most common cause of channel LOF for the mutants examined in this work. Figure S3 further clarifies this correlation by showing that a majority of the mutants exhibiting low current densities are also subject to low cell surface trafficking efficiency.

Because KCNQ1 in cardiac myocytes co-assembles with the KCNE1 protein to form a complex early in the secretory pathway (*25-27*), we tested whether co-expression of KCNQ1 with KCNE1 alters trafficking. For the 16 mutants tested, co-expression with WT KCNE1 led to, at most, moderate changes in cell surface expression levels (fig. S4). This suggests that for each of the other 35 KCNQ1 mutants, the presence of KCNE1 is likely to exert no more than moderate impact on total channel expression or on surface trafficking, although the possibility of outliers cannot be excluded.

### Proteasomal degradation explains the reduced expression of many KCNQ1 variants

Both low cell surface expression and low surface trafficking efficiencies correlated with low total protein expression (Fig. 1, A and B, table S1). We hypothesized that reduced total expression is due to degradation of nascent KCNQ1 via the ERAD/proteasomal pathway (*28*). To test this, cells transfected with poorly expressing KCNQ1 mutants were treated for 20 hours with a proteasome inhibitor, MG132 (25 μM). As seen in Fig. 2, proteasomal inhibition resulted in only modest changes in surface expression, but usually resulted in dramatically enhanced total expression levels for these mutants, although not for WT. These results implicate the endoplasmic reticulum protein folding quality control system and the coupled ERAD-to-proteasome pathway as the major determinant of the differences in total expression levels between WT and most trafficking-deficient KCNQ1 mutants. Mistrafficking-prone mutants are more efficiently targeted for degradation than WT and variants that traffic with near-WT efficiency.

**Figure 2.**
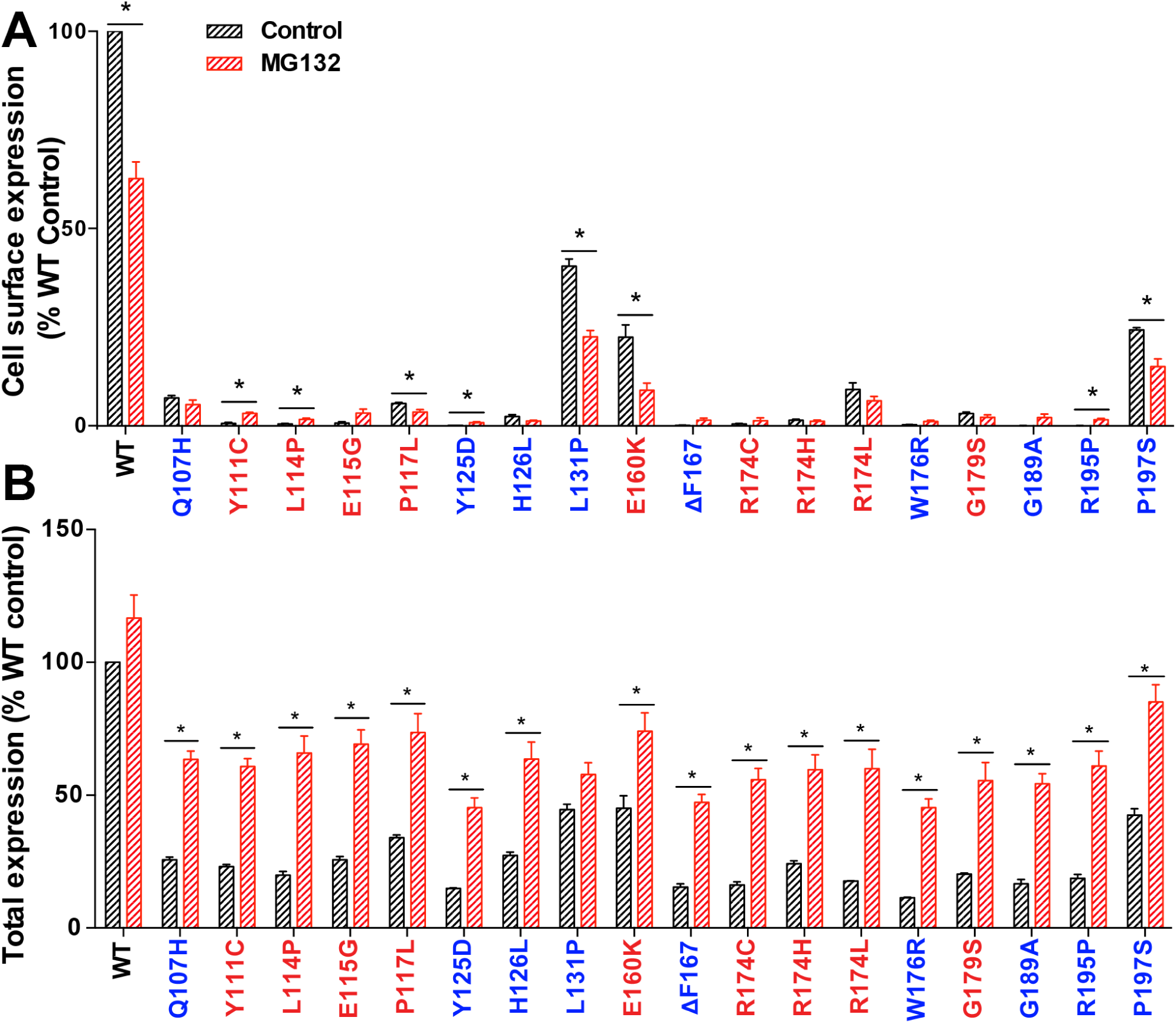
Treatment of cells with a proteasome inhibitor (MG132) has modest impact on surface expression levels of the trafficking-deficient KCNQ1 variants but often increases the total expression. Cells expressing WT or mutant KCNQ1 were treated with 25 μM MG132 or vehicle for 20 h. Cells were fixed and permeabilized (this step was omitted for measuring the surface expression) and then stained with myc-tag mouse monoclonal antibody. Cells were then washed and stained with anti-mouse Alexa Fluor 647 antibody. Immunostaining of 2500 cells was quantitated by flow cytometry. Results are expressed as mean ± S.E.M. for at least three independent experiments. Data are colored coded: LQTS mutant (red) VUS (blue) or predicted neutral polymorphism (black). Data labeled with an asterisk and a horizontal bar indicate those for which the measured KCNQ1 protein level for vehicle-treated cells was statistically different from the level measured in MG132-treated cells (P<0.05).

### Impact of co-expressing mutant and WT KCNQ1 on cell surface channel levels

Because of the dominant nature of type 1 LQTS and the heterozygous (WT/mutant) state of KCNQ1 patients, we examined the effect of co-expressing trafficking-defective mutants with WT channels on expression levels. As shown in Fig. 3, most of the tested mutants exert a partially dominant-negative effect on WT surface expression, with this trend correlating with LOF. For example, most of the mutants that exhibit non-detectable cell surface expression when expressed alone (*e.g.* G189A and R195P) exhibited WT/mutant cell surface expression levels that are well below 40% of WT/WT (Fig. 3). These results are consistent with some degree of dominant-negative suppression of WT trafficking by mutant KCNQ1. Some trafficking-impaired mutants appear to be especially effective at disrupting WT surface expression. For example, surface expression of WT/L114P is below 10% of WT/WT, suggesting that the presence of even a single L114P subunit within the WT/L114P heterotetrameric channel is sufficient to impair trafficking of the entire complex. At the other end of the spectrum, co-expression of WT with L236R appears to restore cell surface expression of L236R. Between these extremes are cases such as WT/H126L, where surface expression is roughly 40% of WT/WT levels. For such cases a possible explanation is that heterotetramers containing one or even two H126L subunits may pass ER quality control and traffic to the cell surface. However, it is not possible to exclude an alternate explanation that mutants such as H126L are so folding defective that they cannot co-assemble with WT KCNQ1, such that only WT-only homotetramers form.

**Figure 3.**
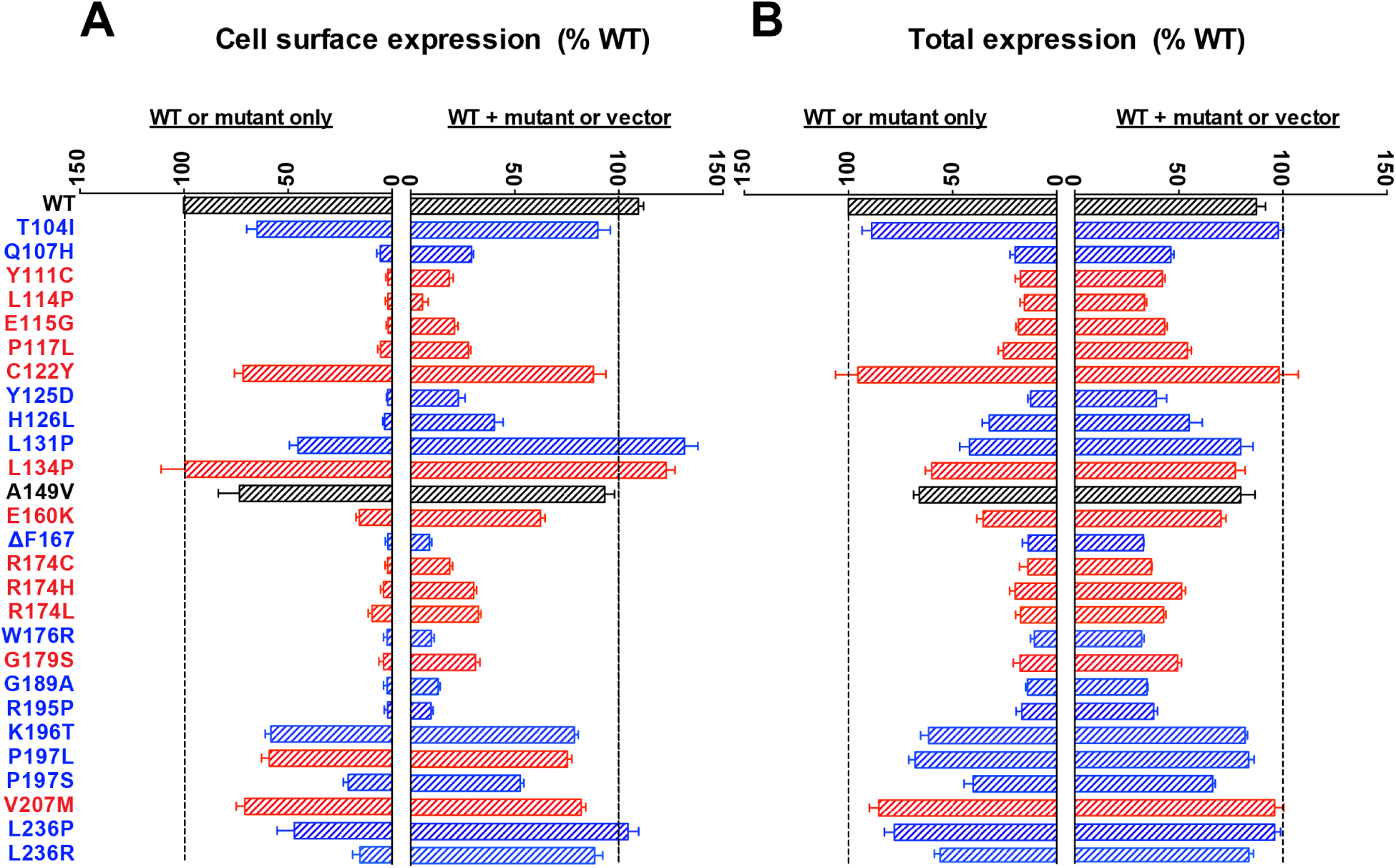
Effect on KCNQ1 trafficking of co-expression of WT KCNQ1 with mutant KCNQ1. (**A**) Total WT+mutant expression levels and (**B**) surface WT+mutant expression levels. HEK293 cells were transiently transfected either with O.5 μg WT or mutant plasmid only (results on the left of each panel) or were co-transfected with both O.25 μg WT and O.25 μg mutant plasmids (heterozygous conditions, results presented on the right of each panel). See the legend of Fig. 1 for additional details.

Overall, most WT/mutant pairs exhibit cell surface expression levels intermediate between those observed for mutant-only and WT-only conditions. Given that most of the mutation sites are located in parts of the VSD that do not make contacts with other subunits, the simplest model to explain the varied impact of mutant coexpression on trafficking is that WT and most mutants coassemble to form tetramers of all possible WT:mutant stoichiometries, with mutant-only and WT-only homotetramers being the least populated states. WT/mutant heterotetramers will exhibit a range of trafficking efficiencies determined by the ratio of WT/mutant subunits. These results indicate that while mutant-only expression assays are informative and yield data that correlates well with functional measurements (as in Fig. 1), additional insight can be gained by also conducting experiments in heterozygous conditions.

### Use of NMR spectroscopy to identify severely folding-destabilized KCNQ1 variants

In an attempt to gain mechanistic insight into how mutations in KCNQ1 altered channel trafficking and function we collected 2D ^1^H,^15^N-TROSY NMR spectra for WT and mutant forms of the VSD. TROSY spectra exhibit contour peaks for each amide ^1^H/^15^N pair along the protein backbone and provide a “fingerprint” pattern that yields insight into mutation-induced changes in protein structure and/or stability (*29*). The WT KCNQ1 VSD yields a well-resolved spectrum (Fig. 4A) in lyso-myristoylphosphatidylglycerol (LMPG) micelles, consistent with the protein being well-folded. This spectrum was used as a reference for comparison with the spectra from 47 of the mutants examined in this work. Spectra were not acquired for the severely-mistrafficking L114P, ΔF167, W176R, and R195P mutants because they failed to express to a sufficient level in *E. coli* for preparation of an NMR sample, consistent with the hypothesis that they are severely unstable and/or misfolded, which is supported by additional data below.

**Figure 4.**
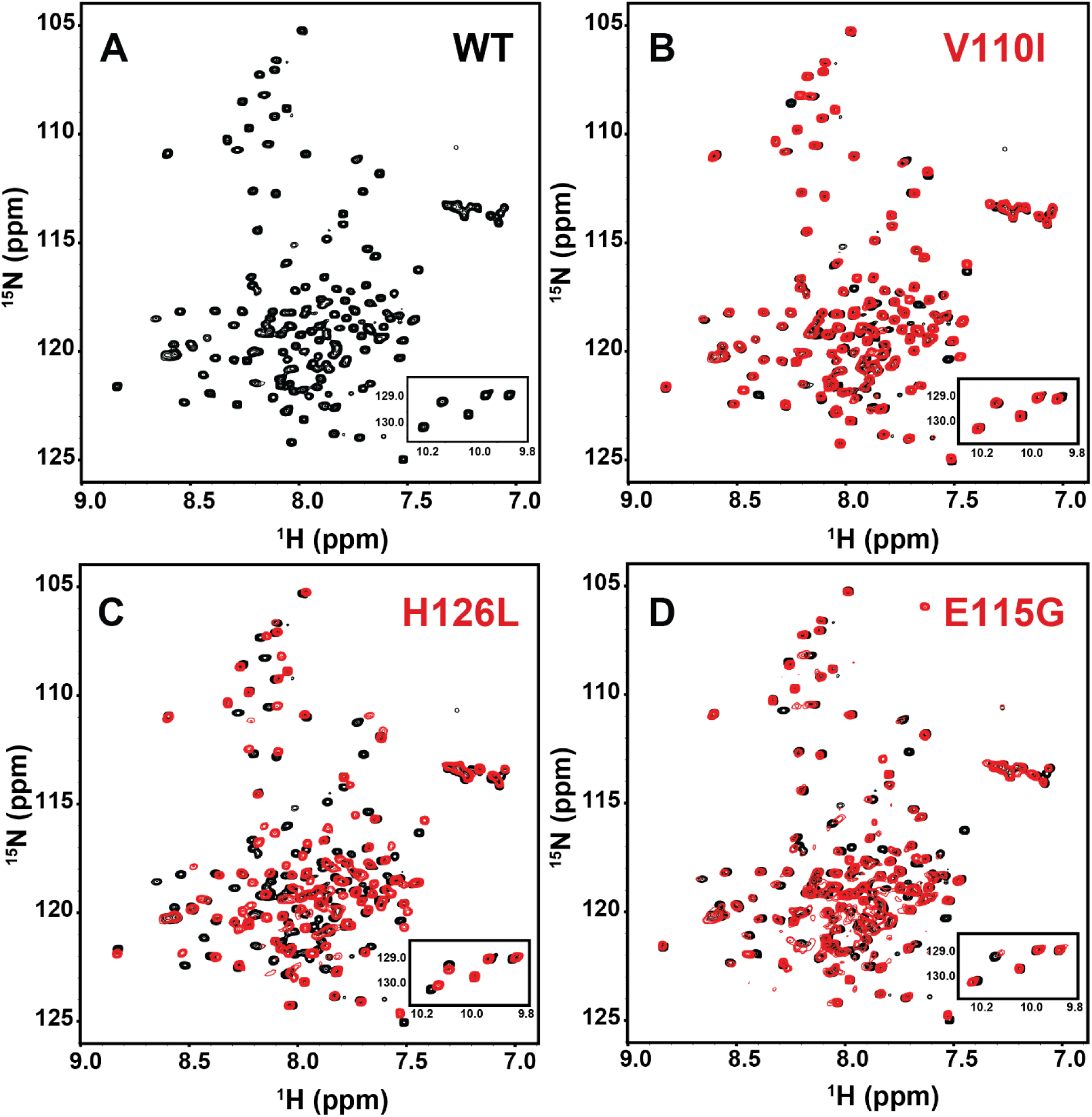
9OO MHz ^1^H,^15^N-TROSY NMR spectra of the WT KCNQ1 VSD (residues 1OO-249) (A) and representative mutant forms (**B-D**). The spectrum of each mutant VSD is shown in red, superimposed on the black spectrum of the WT VSD spectrum. Data were collected at 5O°C for WT and mutant forms of VSD in LMPG micelles at pH 5.5. The LMPG concentration for all samples was 5O-8O mM and the KCNQ1-VSD concentration was O.3 mM in all samples. Panel (**B**) illustrates the spectrum from the V11OI mutant, for which the only changes relative to the WT spectrum are shifts in peak positions. Panel (**C**) illustrates the spectrum from the H126L mutant that is deemed to be moderately destabilized based on a modest degree of line broadening for a number of peaks relative to the corresponding peaks in the WT spectrum. Panel (**D**) illustrates the spectrum from the E115G mutant, which is deemed to be severely destabilized based on extensive peak broadening and even disappearance of a number of peaks.

Among the 47 mutants examined by NMR (Fig. 4, B to D and fig. S5), two major classes of spectral changes were observed relative to the WT spectrum. First, all spectra exhibited some changes in peak positions (c.f. Fig. 4B). These include differences for backbone amide peaks representing residues at or proximal to the mutation site. Shifts in peak positions for amides at some sequentially distal sites were also usually seen, which likely reflect minor distortion of tertiary structure and/or dynamics. There were no overt correlations between shifts in peak positions and either channel function or cell surface trafficking. Assessing the number of peaks that shift or the shift magnitudes also did not correlate with function and trafficking. This is not surprising given that ^1^H and ^15^N NMR chemical shifts are sensitive to many different factors and cannot readily be interpreted in conformational terms. We also found no evidence that shifts in peak position reflect a change in the equilibrium between two (possibly active and inactive) VSD states. We conclude that the differences in amide ^1^H-^15^N peak *positions* between these mutants and WT are not informative.

A second class of NMR spectral differences was observed for a subset of 13 mutants. These mutants yielded spectra in which many of the peaks were significantly *broadened*, in some cases to the point of complete disappearance. Examples are shown for moderate (Fig. 4C) and severe (Fig. 4D) cases (see also fig. S5 and table S1). Extensive mutation-induced peak broadening in ^1^H,^15^N-TROSY or HSQC NMR spectra is a hallmark of incompletely folded “molten globular state” proteins (*30*). This phenomenon is the outcome of mutation-induced destabilization of the folded state to an extent where the most highly populated state is that of a folding intermediate in which multiple conformations exchange with each other and/or with the folded and unfolded states at an intermediate rate on the NMR time scale. Such “intermediate exchange” results in extensive NMR line broadening. The spectra of these 13 KCNQ1 mutants establish that the structures of these mutants are significantly destabilized relative to WT. The fact that this phenomenon is more severe for some mutants (see Fig. 4D) than others (e.g. Fig. 4C) serves to identify the mutants that are most severely destabilized. It is notable that the 13 mutants showing moderate-to-severe peak broadening also exhibited loss of channel function and low total and surface expression levels (in all cases <65% of WT values, see table S1). For these mutants, disease-associated LOF appears to stem from mutation-induced destabilization of the protein, resulting in impaired trafficking to the cell surface and enhanced degradation of the protein by the ERAD/proteasomal pathway, a pathway it shares with HERG (*31*) and some other channels(*32*). These 13 mutants also tended to express more poorly in *E. coli* than trafficking-proficient mutants. Based on this it is reasonable to assume that the 4 mutants that completely failed to express in *E. coli* are also severely folding-destabilized.

### S0 is a central structural and dynamical element of the VSD

Five of the LQTS mutants in the little-characterized S0 segment were seen by NMR to be folding-defective (Q107, Y111C, L114P, E115G, and P117L) (red side chains in Fig. 5A). These and other LQTS sites in S0 directly interact with seven LQTS or VUS sites in S1 and S2 for which mutants also are mistrafficking-prone (Y125D, H126L, R174C, R174H, R174L, W176R, and G179S; magenta side chains in Fig. 5A). To further probe the interactions of S0 with the rest of the VSD we conducted molecular dynamics (MD) simulations on the VSD in an explicit 37°C dimyristoylphosphatidylcholine (DMPC) bilayer. Three independent 500 nsec MD runs were carried out, for a total of 1.5 μs of simulation time.

**Figure 5.**
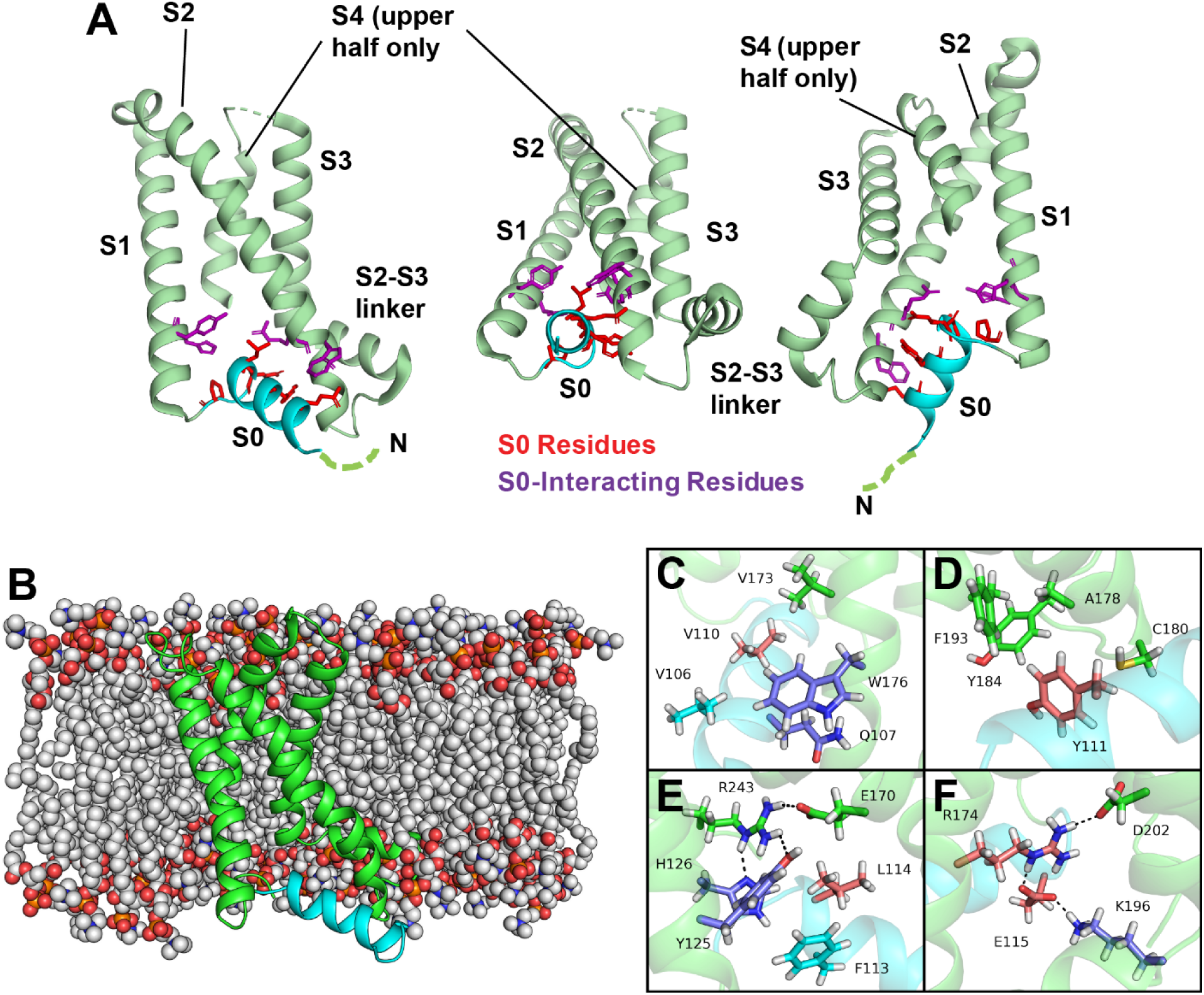
Structural locations and key interactions involving mutations sites and/or SO in the KCNQ1 VSD. (**A**) Three views of the VSD illustrating the locations of the 5 sites in SO (red side chains: Q1O7, Y111, L114, E115, and P117) subject to LQTS mutations that destabilize the VSD, resulting in mistrafficking and degradation of KCNQ1. Shown in magenta are the side chains for four residues that interact with these SO residues and that are also subject to destabilizing VUS or LQTS mutations resulting in channel LOF. The open state VSD coordinates from the cryo-EM structure of KCNQ1 (Protein Databank ID: 5VMS) were used to generate this figure (*23*). The V11O LQTS mutation site is also located in SO, but the mutation does not appear to destabilize the protein. Panels **B-F** illustrate results from the MD simulation of the KCNQ1 VSD. (**B**) Structural model of the WT open state human VSD in a DMPC bilayer after 500 ns of MD. The VSD is displayed in cartoon representation with S1-S4 colored pale green and the S0 helix colored cyan. DMPC molecules are depicted as spheres and colored by atom identity: C – grey, O – red, N – blue, P – orange. (**C – F**) Non-bonded interactions involving sites in S0 and sites contacting S0 for which LQTS and VUS mutations were characterized in this study (see Results). Amino acid side chains are drawn as sticks. LQTS and VUS mutation sites are colored light red and blue, respectively. Residues for which mutations are neutral or have not been characterized in this work, are colored grey and green, respectively. Predicted hydrogen bond interactions are indicated by black dotted lines and atoms. The nature of the non-bonded interactions involving S0 is further described in the main body of the text and in table S2.

In each of these calculations, the VSD maintained its original structure and reached equilibrium after ca. 50 ns of simulation time (see fig. S6A). Analysis of the trajectories revealed a dense network of interactions of S0 with other parts of the VSD, as well as with water and lipids (Fig. 5B and fig. S6B). Important observed contacts that involve the LQTS and VUS sites are listed in table S2. The N-terminal half of S0 is bounded by the C-terminal end of S2 and involves π-π stacking between VUS mutation sites Q107 (S0) and W176 (S2) (Fig. 5C). The indole side chain of W176 also interacts with S0 hydrophobic residues V106 and V110 on one side and is exposed to lipid on the other side. The W176R mutation would eliminate the π-π interaction and also introduce a charged side chain adjacent to lipid. Similarly, residues in S2 (A178, C180) and the S2-S3 linker (Y184, F193) form a hydrophobic pocket to accommodate the phenol side chain of S0 LQTS site Y111 (Fig. 5D). Long-range electrostatic interactions are also possible between Y111 and the side chains of R174, R181 and K196. This suggests that the Y111C LQTS mutation weakens these hydrophobic and electrostatic interactions, accounting for the severe destabilization of this mutant as documented by NMR (fig. S5 and table S1).

In the C-terminal half of S0, LQTS site L114 makes hydrophobic contacts with Y125 (S1) and V173 (S2) (Fig. 5E), and polar interactions are seen between its backbone CO and side chains of H126 (S1), R243 (S4) and Q244 (S4). The S0 L114P LQTS mutation, which led to a significant reduction of protein expression in HEK293 cells and failure to express in *E. coli* would be expected to perturb not only tertiary interactions but the secondary structure of the S0 helix itself and its interaction with the rest of the VSD, thereby disrupting the coordinated network of interactions S0 makes with various sites in S1, S2, the S2-S3 linker and S4. Two of the L114-interacting residues, Y125 and H126, are sites in S1 that are subject to VUS mutations resulting in loss of function and mistrafficking (Fig. 5). These residues bridge S0 with sites in S4 known to be critical for voltage-sensing—H240, R243, and Q244 (*2*) (Fig. 5E)—and therefore could be important not only for the stability, but also in the voltage-sensing function of the VSD. The carboxyl chain of S0 LQTS mutation site E115 was seen to frequently form hydrogen bonds with R174 in S2 and K196 in the S2-S3 linker (Fig. 5F) and also forms transient hydrogen bonds with the R243 and R249 in the S4 helix. R174 is an LQTS site and seems to be central to a network of hydrogen bonds that spans all transmembrane helices, connecting sites in S0 (E115) with sites in S2, S3 (D202), and S4 (R243, Q244).

Finally, S0 and the preceding N-terminal segment were also seen to interact with surrounding membrane lipids and water (see fig. S7). Arginine residues R103 and R109 participated in frequent hydrogen bonding interactions with phosphodiesters of DMPC. These protein-lipid interactions likely help to anchor the VSD in the membrane and thus represent another mechanism by which S0 contributes to the stability of the VSD.

## Discussion

### Classifying mechanisms of KCNQ1 loss-of-function

We recently completed a high-throughput electrophysiological investigation of KCNQ1 variants including the 51 mutants studied in this current work (*22*). Although the focus of the previous study was to illustrate a new and efficient paradigm for linking genotype to function for human ion channels, observations made about the functional consequences of the variants were informative. In particular, a large number of variants (32 of 51) exhibit substantially lower (e.g., <65%) peak current density than WT channels consistent with loss-of-function. Four variants trafficked normally and exhibited normal peak channel currents but were deemed to be dysfunctional because of major alterations in channel V1/2 for activation or perturbed deactivation kinetics. Other variants, including 11 out of 13 predicted neutral amino acid substitutions exhibited normal or near normal function. Among variants associated with LQTS, most (∼85%) exhibited loss-of-function traits, but the mechanisms responsible for this functional impairment was not further explored. In this paper we sought to elucidate the mechanisms of channel LOF by quantifying total and cell surface expression levels for WT and mutant forms of KCNQ1.

We observed that 23 of 32 LOF mutants exhibited lower cell surface expression (≤65%) than WT. Further analysis leads to the conclusion that each KCNQ1 mutant examined in this work can be classified into one of 6 categories, as summarized below and listed in Table 1.

**Table 1.** Classification of KCNQ1 variants

| **Classification<sup>1</sup>** | **Criteria** | **Variants<sup>2</sup>** |
| --- | --- | --- |
| I | Normal expression/trafficking, but low peak current density. Dysfunctional channel. | V110I C122Y L134P A150T A150V T169M K218E I227L Q234P |
| II | Normal channel function, but low expression and/or trafficking efficiency. | P197L |
| III | Normal surface levels and peak current density, but altered channel $V_{1/2}$ and/or deactivation rate. | H105L<sup>3</sup> R109L T118S I132L |
| IV | Defective in both channel properties and expression/trafficking. | T104I L131P E160K K196T P197S L236P L236R |
| V | Severely trafficking-defective. Current is so low that channel properties cannot be assessed. | Q107H Y111C L114P E115G P117L Y125D H126L ΔF167 R174C R174H R174L W176R G179S G189A R195P |
| VI<br>(WT-Like) | WT-like properties in all tested | V100I A102S T104S H105N H105Y V106I V124I F127L A128T V129I V133I V135A V135I A149V V207M |

<sup>1</sup> The 32 mutants in classes I, II, IV, and V are all deemed to be loss of function because they exhibit maximal channel conductance $\leq 65\%$ of WT. The 4 class III mutants exhibit normal maximal peak currents, but altered $V_{1/2}$ and/or deactivation rates, and are therefore deemed dysfunctional.
<sup>2</sup>The listed mutants are color coded to indicate their initial classifications prior to this work as LQTS (ref), VUS (cyan), or neutral/benign.
<sup>3</sup>H105L exhibited a hyperpolarizing shift in $V_{1/2}$ for channel activation, R109L exhibited a hyperpolarizing shift in $V_{1/2}$ for activation, T118S exhibited a depolarizing shift in $V_{1/2}$ for activation, and I132L exhibited slowed deactivation.

*Class I mutants exhibit normal cell surface expression (≤65% that of WT), but exhibit reduced channel conductance.* The mechanism for LOF induced by each such mutation is reduction of channel conductance within otherwise normally folded and trafficked protein. Nine of 32 LOF mutants belong to this class.

*Class II mutants have normal channel properties when properly folded and surface-trafficked, but exhibit dramatically reduced cell-surface expression (only 10-65% of WT)*. Only 1 variant (P197L) belongs to this class.

*Class III mutants reach the cell surface at near normal levels and display WT-like channel conductance, but exhibit significant alterations in the voltage-dependence of channel activation and/or altered deactivation kinetics, indicating dysfunctional channel properties.* Four mutants fell into this category.

*Class IV mutants exhibit low cell surface expression (10-65% of WT) with LOF being compounded by the fact that even the population of channels that reaches the cell surface is dysfunctional.* Seven mutants belong to this class.

*Class V mutants exhibit cell surface levels that are <10% that of WT.* The plasma membrane levels of these mutants are so low that it is not possible to accurately determine whether the very small population of assembled and trafficked channel is also dysfunctional. Fifteen mutants belong to this class.

*Class VI mutants exhibit WT-like channel properties, total expression and surface trafficking.* Fifteen of 51 mutants exhibited normal channel trafficking and function. These included 11 out of 13 of the predicted benign mutant forms. The other two, T118S and A150V, exhibited properties that caused them to be classified just outside of the range of “normal”. That these two predicted benign mutants exhibited significant dysfunction may point to the peril of assuming that sequence variation at a particular protein site between orthologs represents neutral evolutionary drift. The exact functional properties required for the KCNQ1/KCNE1 I_Ks_ channel in the regulation of human heartbeat may be significantly different from the orthologous channel complex in other organisms, such that sequence variation may reflect functionally essential adaptations. Conversely, it must be pointed out that 3 LQTS mutants exhibited WT-like properties, meaning either that these mutants have defects that we failed to uncover or that they are misclassified.

It seems possible that the personalized treatment of LQTS may someday benefit from ascertaining which channel LOF mechanism pertains to a given patient. For example, the appropriate therapeutic approach for a LQTS patient with KCNQ1 that traffics normally, but has defective channel properties is likely to be different that for a patient with KCNQ1 that is folding-destabilized, leading to mistrafficking and degradation. The tailoring of therapies to varying classes of channel defects has become a reality for certain cystic fibrosis transmembrane regulator (CFTR) mutations that cause cystic fibrosis, and may soon become applicable to other disease-membrane protein relationships. For cystic fibrosis, the most common disease-causative mutation is ΔF508 in the CFTR chloride channel. ΔF508 CFTR does not normally traffic to the plasma membrane but, when coaxed to do so by pharmacological chaperones, is then seen to be partially active (*33-35*). Many others of the hundreds of known CFTR mutants are likely to be trafficking-defective and potentially rescuable by a pharmacological approach (*36*). On the other hand, some CFTR mutants are known to traffic normally, but have dysfunctional channel properties.

Drugs have been developed that specifically improve the functionality of some such mutants (*33, 34*). Whether this will prove feasible for KCNQ1 remains to be seen.

### KCNQ1 misfolding caused by underlying instability is a common LOF mechanism

We observed that the majority (23/32) of the LOF mutants examined in this work exhibited much lower levels of cell surface expression than WT. This result is reminiscent of the conclusion from studies of the HERG potassium channel that most type 2 LQTS disease mutations in this protein result in trafficking defects (*31*). These KCNQ1 mutants may also be analogous to Charcot-Marie-Tooth disease (CMTD) mutant forms of the human tetraspan membrane protein peripheral myelin protein 22 (PMP22). Most CMTD mutant forms of PMP22 traffic to the plasma membrane with lower efficiencies than WT PMP22 (*37*), and it was shown that the efficiency of trafficking correlates linearly with disease severity (*38*). Moreover, a linear relationship was also observed between the magnitude of energetic destabilization of PMP22 structure and both its intracellular retention (mistrafficking) and disease severity (quantified as patient nerve conductance velocity) (*38*). That study established mutation-induced destabilization of folded PMP22 as the defect underlying many phenotypes of CMTD.

In the present work, we have examined whether the same relationship holds between mutation-induced folding-destabilization of KCNQ1 and LQTS. Mutants have been identified for which structural destabilization is detectable by NMR. Out of 23 KCNQ1 mistrafficking-prone mutants, 17 were determined to be structurally unstable. It is feasible that the remaining 5 mistrafficking mutants are also folding-destabilized, just not to an extent that is evident in their NMR spectra. Significantly, none of the 18 mutants characterized in this work exhibiting WT-like function and trafficking displayed NMR spectral traits that were very different from WT. From these results we conclude that, at least for 32 KCNQ1 LOF mutants examined herein, the most common defect that results in loss of channel function and LQTS is mutation-induced destabilization of the protein, leading to retention by endoplasmic reticulum protein folding quality control and ERAD pathway-based degradation by the proteasome. Mistrafficking and degradation via the ERAD pathway has previously been documented for numerous mutants forms of the HERG channel that result in type 2 LQTS (*31*). Whether destabilization of KCNQ1 structure will prove to be the most common mechanism underlying LOF for the hundreds of known LQTS mutant forms of KCNQ1 remains to be seen. One also wonders whether drug-like small molecules might be developed that can reach and bind nascent KCNQ1 in the endoplasmic reticulum to stabilize otherwise misfolding-prone channel mutants, leading to restoration of trafficking to the cell surface and possibly channel function.

PMP22 and KCNQ1 are currently the only two human disease-linked membrane proteins for which the relationship of protein stability to disease mechanism has been definitively established for more than a few select mutations (such as the well-characterized ΔF508 form of the CFTR channel). However, given that many other diseases are linked to mutation-induced membrane protein misfolding, such as retinitis pigmentosa (rhodopsin), Pelizaeus-Merzbacher disease (proteolipid protein}, cystic fibrosis (CFTR), and diabetes insipidus (vasopressin V2 receptor), it is likely that mutation-induced destabilization is also a trigger for the disease-causative intracellular retention and/or degradation of these proteins. There is a pressing need to develop general methods to quantitatively compare the stabilities of WT and disease mutant forms of these proteins. It is also interesting to note that some of the mutation sites associated with destabilization of KCNQ1 correspond via homology to known disease mutation sites in other voltage-gated ion channels (*39-41*), including a number of KCNQ2 and KCNQ3 sites for which mutations are associated with benign familial neonatal seizures and epileptic encephalopathy (see http://www.hgmd.cf.ac.uk). This suggests that mutation-induced destabilization of protein folding is likely a very common underlying mechanism for many other channelopathies.

### The S0 helix stabilizes the KCNQ1 VSD

The S0 segment found in many voltage-gated channels has previously received little attention as to its roles in VSD structure and function. (*Note that the S0 segment in KCNQ1 and other Shaker-type channels should not be confused with the transmembrane S0 helix in BK channels*). Our results reveal a critical role for this widely-conserved structural element. Of the 17 KCNQ1 mutants found in this work to be significantly folding-destabilized, 5 involve LQTS mutations at sites located in S0 (Q107H, Y111C, L114P, E115G, and P117L; see fig. S1 and Fig. 5). We note that S0 mutants Y111C, L114P, and P117L have previously been identified as trafficking-defective (*40, 42*). Another 7 mistrafficking-prone mutants (Y125D, H126L, R174C, R174H, R174L, W176R, and G179S; Fig. 5) involve sites in S1 or S2 that interact with the 5 LQTS sites in S0. MD simulations illuminated the role of S0 as a central scaffolding element that is engaged in a dense network of interactions with other VSD segments, even the functionally critical S4. (Fig. 5). This is further supported by the observation that residues in S0 move in a concerted and correlated fashion with residues in S2 and S4 (see fig. S8). Combined, these observations indicate that S0 contributes to organizing and stabilizing the structure of the VSD.

S0 segments are present in many other voltage-gated channels and even in the VSD-like regulatory domain of transient receptor potential (TRP) channels (*23, 43-47*). As we observed for KCNQ1, disease-causing mutations are sometimes found in the S0 segments of these other channels. For example, each of the 4 distinct S0 segments in the human voltage-gated sodium channel SCN5A (Nay1.5) have at least one known LQTS and/or Brugada syndrome-associated mutation (11 mutations total, see http://www.uniprot.org/uniprot/Q14524 (*23*)). It seems very likely that the critical role for S0 documented in this work for KCNQ1 may extend to other voltage-gated ion channels.

## Conclusions

This study confirms the value of conducting studies of the trafficking and stability of KCNQ1 variants, studies that complement EP functional studies. Not only is significant additional mechanistic information provided regarding the loss of channel function for many variants, but fundamental new insights into KCNQ1 channel structure-function-stability relationships can be gleaned (i.e., recognition of the importance of S0 in VSD structure and stability). This work also highlights the importance of membrane protein destabilization as a potential disease mechanism. The results of this work could conceivably impact future personalized medical decisions for patients with one of the KCNQ1 mutants characterized in this work. Moreover, the results may be used to help train computational algorithms being developed to predict channel functionality and disease outcomes for patients that harbor KCNQ1 variants of unknown significance.

## Materials and Methods

### Cloning

The c-myc tagged human KCNQ1 DNA (GenBank accession number AFOOO571), a gift from Dr. Dan Roden of Vanderbilt University (*24*), was subcloned into a pIRES2-EGFP vector. The c-myc tag (EQKLISEEDL) was introduced into the extracellular S1-S2 linker between Glu146 and Gln147. The VSD, spanning KCNQ1 residues 1OO to 249, was cloned into a pET16b vector with an N-terminal His tag (MHHHHHHG-). Human KCNE1 (L28168) was subcloned into a pcDNA3.1(+) vector. Mutants were generated by QuikChange site-directed mutagenesis using WT *myc-KCNQ1* or *KCNQ1-VSD* cDNA as the template and verified by sequencing to confirm the presence of the desired mutation(s).

### Cell culture and transfection

HEK293 cells were purchased from American Type Culture Collection (Manassas, VA). Cells were cultured in Dulbecco’s modified Eagle’s medium (DMEM) supplemented with 10% fetal bovine serum (FBS), 10 mM HEPES, 1OO units/ml penicillin, and 1OO μg/ml streptomycin at 37^o^C in a humidified atmosphere with 5% CO_2_. HEK293 cells were plated into 6-well plates and transfected with 0.5 μg WT or mutant myc-KCNQ1 per well at 3O-5O% confluence using Fugene 6 transfection reagent (Promega, Madison, WI). When WT and mutant myc-KCNQ1 were co-transfected, the WT/mutant or WT/pIRES2-EGFP vector ratio was 1:1 and total DNA was 0.5 μg. Approximately 48 h later, cells were prepared for flow cytometry measurement. To assess the effect of proteasome inhibitor on myc-KCNQ1 expression, cells were treated starting one day after transfection with 25 μM MG132 or 0.1% DMSO (control) for 20 h before quantitating KCNQ1 protein levels.

### Flow cytometry

On the day of the experiment, cells were placed on ice and washed once with ice-cold PBS containing 25 mM HEPES and 0.1% NaN_3_ (PBS-FC, pH 7.4). Cells were detached in 0.5 mM EDTA in PBS-FC and precipitated by centrifugation at 500 × g for 5 min. As previously described (*38*), cells were then permeabilized and stained using the Fix & Perm kit (ThermoFisher scientific, Waltham, MA) following the manufacturer’s instructions. Briefly, cells were incubated with 100 μl PE-conjugated myc-tag (9B11) mouse monoclonal antibody (Cell Signaling Technology, Danvers, MA) (1:100 dilution in PBS-FC containing 5% FBS) for 30 min in the dark at room temperature. 100 μl fixation medium was then added and cells were incubated for 15 min to be fixed. Cells were then washed once with PBS-FC containing 5% FBS and incubated with Alexa Fluor 647-conjugated myc-tag (9B11) mouse monoclonal antibody (Cell Signaling Technology) (1:100 dilution in permeabilization medium) for 30 min in the dark at room temperature. Cells were rinsed once and fluorescence signals were then measured using a FACS Canto II flow cytometer (BD Bioscience, San Jose, CA). Cells expressing WT myc-KCNQ1 were permeabilized and stained either with PE or Alexa Fluor 647-conjugated antibody to normalize the two fluorescence signals. For the proteasome inhibitor study, cells were fixed and permeabilized (this step was omitted for measuring the surface expression) and then stained with myc-tag (9B11) mouse monoclonal antibody (Cell Signaling Technology) (1:500 dilution in PBS-FC containing 5% FBS) for 30 min at room temperature. Cells were then washed and stained with anti-mouse Alexa Fluor 647 antibody (Cell Signaling Technology) (1:1000 dilution in PBS-FC containing 5% FBS) for 30 min at room temperature in the dark. Single cells expressing myc-KCNQ1 were selected by gating on GFP-positive cells. Fluorescence of cells transfected with the empty vector was used to account for background staining. The expression levels of mutants were calculated as a percentage of the WT channel. The mean and the standard error of the mean (S.E.M.) were calculated from at least three independent experiments. Data were analyzed using GraphPad 6.0 software. The statistical significance of the differences between WT and mutants or between DMSO and MG132 treated cells was determined by Student’s t-test.

### Overexpression and purification of the VSD in *E. coli*

WT and mutant forms of the KCNQ1 VSD were expressed and purified essentially as previously described (*48*). Briefly, KCNQ1 VSD was expressed in Rosetta/C43(DE3) cells and cultured in ^15^N labeled M9 minimal medium. Cells were induced by 1 mM IPTG for 24 h at room temperature and then harvested. Each gram of cells was suspended in 20 ml lysis buffer (75 mM Tris-HCl, 300 mM NaCl, and 0. 2 mM EDTA, pH 7.8) with 5 mM Mg(Ac)_2_, 0.2 mg/ml PMSF, 0.02 mg/ml DNase, 0.02 mg/ml RNase and 0.2 mg/ml lysozyme and tumbled for about 30 min. The lysate was probe-sonicated at 4°C for 5 min with 5 s pulses separated by 5 s. The inclusion body was isolated by centrifugation at 20 kg for 20 min and washed once in lysis buffer. The inclusion body was then solubilized in buffer A (40 mM HEPES, 300 mM NaCl, pH 7.5) containing 0.5% dodecylphosphocholine (DPC) (Anatrace, Maumee, OH) and 2 mM TCEP overnight at 4°C. Insoluble debris was removed by centrifugation at 20 kg for 20 min and the supernatant was incubated with Ni(II)-NTA Superflow resin (Qiagen, Germantown, MD) for at least 1 h at 4°C. The resin was then packed into a gravity-flow column and washed with 10 bed volumes of buffer A containing 0.5% DPC and 2 mM TCEP. Impurities were removed by extensive washing with 12 bed volumes of buffer A containing 0.5% DPC, 2 mM TCEP, and 60 mM imidazole (pH 7.8). DPC on the column was exchanged with LMPG (Anatrace) by washing the column with 10 bed volumes of buffer A containing 0.05% LMPG and 2 mM TECP. The KCNQ1-VSD was then eluted in buffer A containing 0. 2% LMPG, 2 mM TCEP, and 500 mM imidazole (pH 7.8) until A_280 nm_ (as monitored continuously) returned to the baseline level (typically 3 bed volumes).

### NMR spectroscopy

The KCNQ1 VSD (100-249) concentration was determined by A_280_ using an extinction coefficient of 34950 M^-1^ cm^-1^. When needed, more LMPG was added to samples to adjust the ratio of LMPG micelles to protein molecules to 3-5. The eluted protein sample was concentrated 10-fold using an Amicon Ultra centrifugal filter cartridge (30 kDa molecular weight cut-off). The sample was diluted with NMR buffer (50 mM MES, 0.5 mM EDTA, 2 mM TCEP, 0.2 mM LMPG (its critical micelle concentration) pH 5.5) to the initial elution volume and again concentrated 10-fold. The process was repeated a total of three times to ensure efficient buffer exchange. The 200 μl NMR sample containing 0.3 mM KCNQ1-VSD, 50-80 mM LMPG, and 5% D_2_O was then transferred into a 3 mm NMR tube. The 2-D ^1^H,^15^N-TROSY experiment was conducted using the standard Bruker pulse sequence. All NMR data were collected at 5O° C on a Bruker 8OO or 9OO MHz NMR spectrometer.

### Homology modeling of the KCNQ1 VSD structure

A structural model of the KCNQ1 VSD (residues 1OO-249) was generated using the protein structure prediction software package Rosetta (Version 3.8) (*49*) based on the cryo-EM structure of Xenopus KCNQ1 (PDB 5VMS) (*23*) and sequence alignment generated with ClustalW (*50*). 4OOO models of KCNQ1 VSD were assembled through comparative modeling (*51*) using the RosettaMembrane energy function (*52*). Gaps in the threaded model resulting from unresolved regions in the template structure were reconstructed by fragment insertion and cyclic coordinate descent (CCD) loop building (*53*). All models underwent side chain repacking and all-atom refinement while applying a low constraint to the initial coordinates. Models were clustered based on root mean squared deviation (RMSD). The lowest-scoring model of the largest cluster was considered the representative model. Its Cα RMSD compared to the Xenopus KCNQ1 VSD structure was 2.3 Å. MolProbity analysis of the representative model reported an overall score of 1.34 (98th percentile), a clash score of 2.O2 (99th percentile), 14O (95.O%) residues in favored regions of the Ramachandran plot, all residues in allowed regions, 126 (98.4%) favored rotamers, no poor rotamers and no Cβ deviations or bad backbone angles.

### Molecular dynamics simulation of KCNQ1 VSD in a lipid bilayer

A MD simulation of the KCNQ1 VSD was performed in an explicit 1,2-dimyristoyl-sn-glycerol-3-phosphocholine (DMPC) bilayer at 313°K using AMBER16 (*54*) and the LIPID14 force field (*55*). Three independent trajectories, each having a total length of 5OO ns but starting with different input models, were computed. Our ensemble of KCNQ1 VSD homology models was clustered using a hierarchical full-linkage clustering algorithm (*56*) and the centroids of the three largest clusters were chosen as starting coordinates. The coordinates of each input model were aligned with the membrane normal using the PPM webserver (*57*). A complete model of the KCNQ1 VSD in a DMPC bilayer (110 lipids per leaflet) was prepared with the membrane builder tool of the CHARMM-GUI website (*58*). A TIP3P water layer of 20 Å was included, and Cl^−^ ions were added to neutralize the charge of the system. Each bilayer system was first minimized for 15000 steps using steepest descent followed by 15000 steps of conjugate gradient minimization. With the KCNQ1 VSD restrained to its starting coordinates, the lipid and water was heated to 50°K over 1000 steps with a step size of 1 fs using constant boundary conditions and Langevin dynamics with a rapid collision frequency of 10000 ps^-1^. The system was then heated to 100°K over 50000 steps with constant volume dynamics and the collision frequency set to 1000 ps^-1^, and finally to 313°K over 100000 steps with constant pressure dynamics and anisotropic pressure scaling turned on. Positional restraints on the KCNQ1 VSD were then gradually removed over 500 ps, and the system was equilibrated for another 5 ns at 313°K. Production MD was conducted for 500 ns using a step size of 1 fs, constant pressure periodic boundary conditions, anisotropic pressure scaling and Langevin dynamics. MD trajectories were analyzed using CPPTRAJ (Version 15.0) (*59*) and VMD (Version 1.9) (*60*).

## Supplementary Materials

**figure S1** Sequence and structural locations of the human KCNQ1 mutations examined in this work.

**figure S2** Fluorimetric cell flow cytometry assay used to determine total and surface protein expression levels.

**figure S3** Surface trafficking efficiency for each mutant juxtaposed with KCNQ1 peak current density

**figure S4** Effect of co-expression of KCNE1 with KCNQ1 on the surface expression levels of KCNQ1.

**figure S5** ^1^H,^15^N-TROSY NMR spectra (900 MHz) of KCNQ1 mutants (red) superimposed on the spectrum of WT KCNQ1.

**figure S6** Starting vs. ending conformer RMSD vs. time for the MD trajectories and snapshots of the KCNQ1 VSD in bilayers from the MD simulations.

**figure S7** Hydrogen bond-interactions of the KCNQ1 VSD with solvent observed during MD simulations.

**figure S8** Correlation plot for VSD motions from the MD simulations.

**table S1** Functional and trafficking results of 51 human KCNQ1 mutants.

**table S2** List of non-bonded interactions of LQTS and VUS mutation sites in S0 and S0-contacting regions which were observed during MD simulation of the KCNQ1 VSD.

## Acknowledgements

We thank Dr. Brett Kroncke for helpful discussion throughout this project. **Funding:** This work was supported by US NIH grant RO1 HL122010. KCT was supported by NIH fellowship F32 GM117770 and NIH training grant T32 NS00749. GK was supported by a fellowship from the German Research Foundation (KU 3510/1-1). The NMR instrumentation used in this work was supported by NIH S10 RR026677 and NSF DBI-0922862, while the computational resources were supported by NIH S10 OD020154 and NIH S10 RR031634. **Author contributions:** HH, GK, JAS, KCT, AMD, and AH conducted the experiments and calculations of this work. All authors participated in data analysis. HH, CGV, ALG, and CRS wrote the paper with input from all authors. JM, CGV, ALG, and CRS conceived of this work and directed the approaches used. **Competing interests:** The authors declare no competing interests. **Data and materials availability:** All data needed to evaluate the conclusions in the paper are presented in the paper and Supplementary Materials. Materials and additional data are available upon request from the authors. Correspondence and requests for materials should be addressed to C.R.S.

